# Evidence of *Anopheles stephensi* involvement in the transmission of *Plasmodium vivax* in Djibouti, 2024

**DOI:** 10.64898/2026.02.25.707780

**Authors:** Subhika Rao, Jeanne N. Samake, Cristina Rafferty, Peter Mumba, Sheleme Chibsa, Meshesha Balkew, Bouh Abdi Khaireh, Samatar Kayad Guelleh, Mohamed Mousse Ibrahim, Colonel Abdoulilah Ahmed Abdi, Sarah Zohdy

## Abstract

**Purpose:** *Anopheles stephensi* is a malaria mosquito vector that has been raising international concern due to its invasive nature in Africa, including the nation of Djibouti. Since its initial detection in Djibouti in 2012, malaria morbidity and mortality have increased exponentially in the county. While there is an observed association increase in human malaria cases since the arrival of *An. stephensi*, high-quality evidence of *An. stephensi* carrying infective sporozoites is essential to determine the role of the invasive vector in malaria dynamics in Djibouti. This study seeks to confirm the link between *An. stephensi* and malaria transmission in Djibouti and examine genetic relatedness between Djiboutian *An. stephensi* populations and populations across the Horn of Africa. Such information regarding the *An. stephensi* populations and the *Plasmodium* species they transmit is necessary to devise appropriate control strategies and limit malaria transmission within and beyond the country.

**Methods:** One hundred and ninety-six adult *An. stephensi* mosquitoes from Djibouti were collected, molecularly confirmed, analyzed for a portion of the cytochrome c oxidase subunit 1 (COI), and tested for infective sporozoites using a highly sensitive and specific multiplex circumsporozoite enzyme linked immunosorbent assay (csELISA) bead assay. The COI sequences of one hundred and fourteen samples were further used to characterize the population genetic structure of the sampled *An. stephensi* and its genetic relatedness to other *An. stephensi* populations across the Horn of Africa.

**Results:** All 196 samples were morphologically and molecularly confirmed to be *An. stephensi*.

*Plasmodium vivax*210 sporozoites were detected with a positivity rate of 1.02%. An analysis of the COI region showed that the infected *An. stephensi* have the most prevalent COI haplotypes of invasive *An. stephensi* circulating in the Horn of Africa.

**Conclusions:** The findings from this study confirm the involvement of *An. stephensi* in *P. vivax* transmission in Djibouti and describe the genetic relatedness of Djiboutian *An. stephensi* populations to other populations across the Horn of Africa. This highlights the threat of *An. stephensi* invasion and supports a rapid and comprehensive response to mitigate the harm that *An. stephensi* populations cause, particularly through surveillance and control of adult populations.

**Graphical Abstract:** 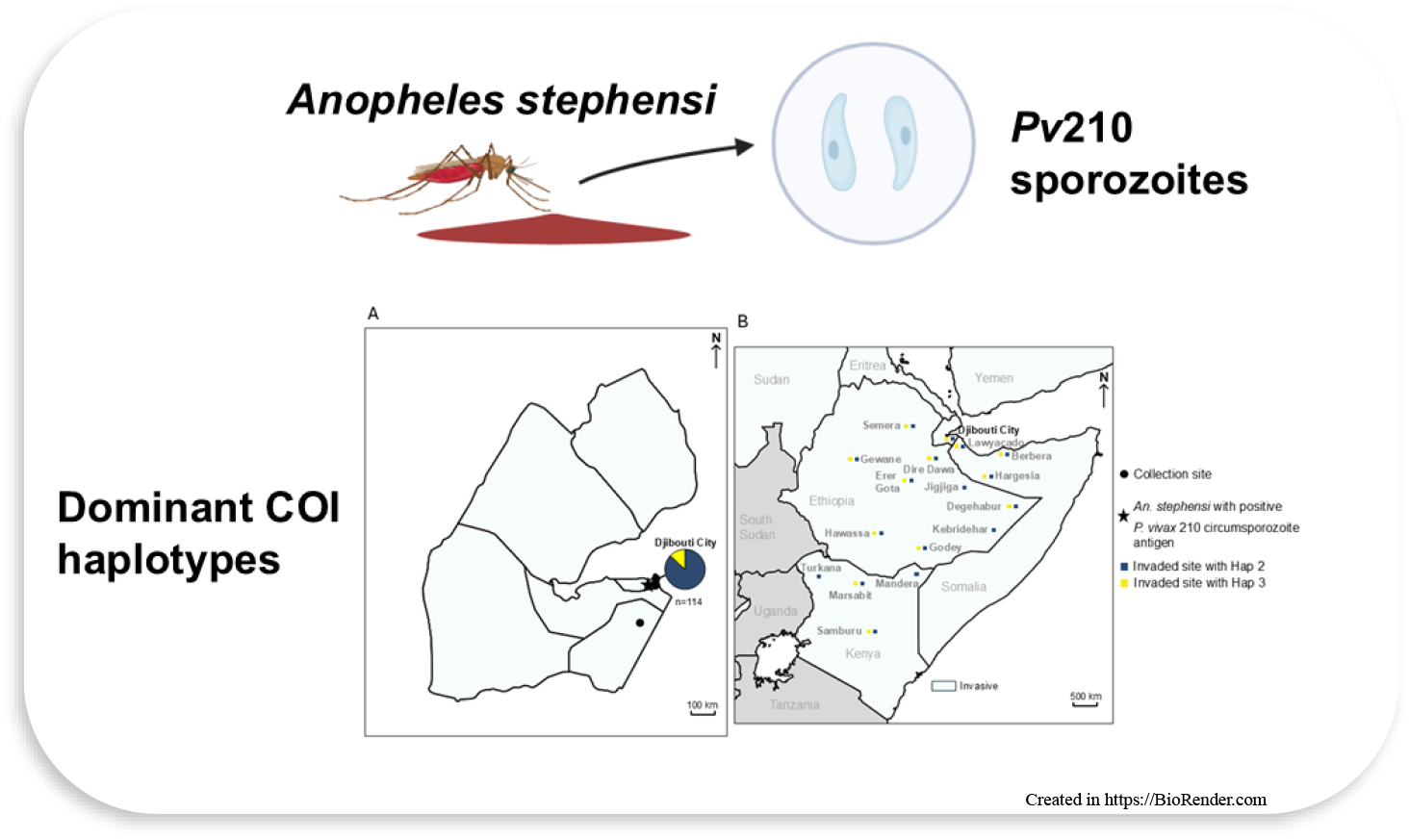

## Background

Malaria presents a significant burden to health systems globally, with 263 million cases and over 597,000 deaths reported in 2023, 94% of which occurs in Sub-Saharan Africa (1). Though there are malaria control programs in place, achieving control and elimination has become increasingly challenging due to emerging threats such as insecticide resistance and invasive malaria vectors like *Anopheles stephensi*.

Native to South Asia and parts of the Middle East, and capable of transmitting both *Plasmodium falciparum* and *P. vivax, An*. s*tephensi* was first detected in Africa in Djibouti in 2012 (2). In subsequent years it was detected in multiple African countries, including Ethiopia (2016), Sudan (2016), Somalia (2019), Nigeria (2020), Ghana (2022), Kenya (2022), Eritrea (2022), and Niger (2024) (3).

In Djibouti, since the first detection of *An. stephensi* in 2012, malaria cases have increased more than 20-fold, from 1684 cases in 2013 to 38,944 cases in 2023 (1). The unprecedented increase includes both *P. falciparum* and *P. vivax* malaria cases (1). This is highly concerning considering Djibouti was nearing the malaria pre-elimination phase prior to 2012 (4). Also, unlike native African malaria vectors, *An. stephensi* persists year-round due to its ability to breed in artificial containers in both urban and peri-urban settings (5-7). Indeed, *An. stephensi* has surpassed the native African malaria vector *An. arabiensis* to become the primary vector in Djibouti City (7, 8). It was found implicated in a resurgence of malaria cases in Djibouti City in 2014 (2) and 2019 (9) through positive infectivity for both *P. falciparum* and *P. vivax* based on a qualitative dipstick assay (10). *Anopheles stephensi* was also suspected in an outbreak among French military personnel in Djibouti City in 2019 (7).

However, the first direct link of the invasive *An. stephensi* involvement in malaria transmission was reported during a malaria outbreak in an urban setting and during the dry season in the neighboring country, Ethiopia, in 2022 (11). A multiplex bead assay of the circumsporozoite enzyme-linked immunosorbent assay (csELISA) (12) was used to molecularly confirm the involvement of *An. stephensi* in malaria transmission in Ethiopia. Thus, in this study, we aim to (1) use a similar molecular approach to determine whether *An. stephensi* is linked to the ongoing malaria outbreak in Djibouti City, and (2) determine its relatedness to other *An. stephensi* in the Horn of Africa. Understanding the role *An. stephensi* plays in the high rates of malaria transmission in the country will inform targeted vector control approaches and surveillance of this invasive malaria species in Djibouti and the invaded region in Africa.

## Methods

Adult *Anopheles* mosquitoes were collected in February 2024 through backpack aspiration in Djibouti City, the capital city of Djibouti, where over 90% of malaria cases have occurred in recent years (13). All 196 samples collected were morphologically identified as *An. stephensi*, molecularly confirmed, then analyzed for a portion of the cytochrome c oxidase subunit 1 (COI) using previously described protocols (14). All 196 samples were then tested for *Plasmodium* infectivity using the highly sensitive and specific multiplex csELISA bead assay as previously described (11, 12). Briefly, mosquito heads-thoraces were separated from legs, wings, and abdomens between the second and third legs from all 196 samples, homogenized and tested for *P. falciparum, P. vivax*210, and *P. vivax*247 following the csELISA protocol detailed in Sutcliffe et al., 2021(12).

Furthermore, the COI DNA sequences were analyzed to generate population genetic statistics and conduct phylogenetic analysis to determine the level of genetic diversity, population structure of the sampled *An. stephensi*, and relatedness to other *An. stephensi* populations in the Horn of Africa. Briefly, basic statistics were generated in DNAsp version 6 (15), including the number of polymorphic (segregating) sites (s), number of haplotypes (h), haplotype diversity (Hd), and nucleotide diversity (π). Next, the COI sequences were further aligned in CodonCode with previously published *An. stephensi* COI sequences from the Horn of Africa retrieved from Genbank. Phylogenetic relationships were inferred using a maximum likelihood approach with RAxML GUI using the GTR model of nucleotide substitutions, gamma model for rate of heterogeneity (GTRGAMMA), and one thousand replicates in one run for bootstrap analysis (16). The tree with the highest log likelihood was visualized and formatted using FigTree v1.44 (17). Lastly, a Horn of Africa map showing the distribution of *An. stephensi* most prevalent COI haplotypes (i.e., Hap 2, and Hap 3) was generated including data from Carter et al. 2021(18), Samake et al. 2024 (5), and Hawaria et al. 2023 (19) (Ethiopia), Ali et al. 2022 (20) (Somalia), and Samake et al. 2025 (21) (Kenya).

## Results and Discussion

Two out of the 196 samples (1.02%) tested positive for the presence of the *Pv*210 circumsporozoite protein, which indicates the presence of malaria in Djiboutian *An. stephensi* (Table 1). The genetic diversity statistics based on 114 out of the 196 generated COI DNA sequences revealed two segregating sites leading to two haplotypes (Table 2). Based on the phylogenetic analysis, the two haplotypes clustered with previously reported *An. stephensi* COI haplotypes in Djibouti, namely, haplotype 2 (n=99) and haplotype 3 (n=15) (22) (Figure S1). These two haplotypes were reported as the most widespread COI haplotypes of the invasive *An. stephensi* in Djibouti and across the Horn of Africa (18, 22) (Figure 1).

**Table 1.**
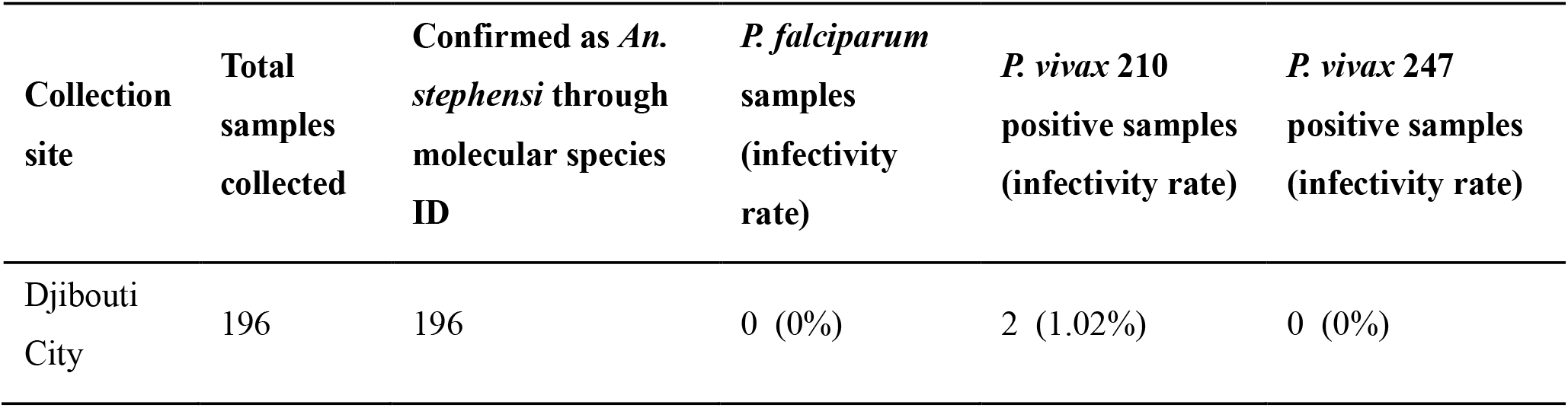
Sample information and infectivity rate.

| Collection site | Total samples collected | Confirmed as *An. stephensi* through molecular species ID | *P. falciparum* samples (infectivity rate) | *P. vivax* 210 positive samples (infectivity rate) | *P. vivax* 247 positive samples (infectivity rate) |
| --- | --- | --- | --- | --- | --- |
| Djibouti City | 196 | 196 | 0 (0%) | 2 (1.02%) | 0 (0%) |

**Table 2.** Population genetic diversity based on mtDNA COI loci of *Anopheles stephensi* from Djibouti, 2024.

| Collection site | Sample size (N) | Number of polymorphic (segregating) sites (S) | Number of haplotypes (h) | Haplotype diversity, (Hd) | Nucleotide diversity ($\pi$) |
| --- | --- | --- | --- | --- | --- |
| Djibouti City | 114 | 2 | 2 | 0.231 | 0.00145 |

**Figure 1.**
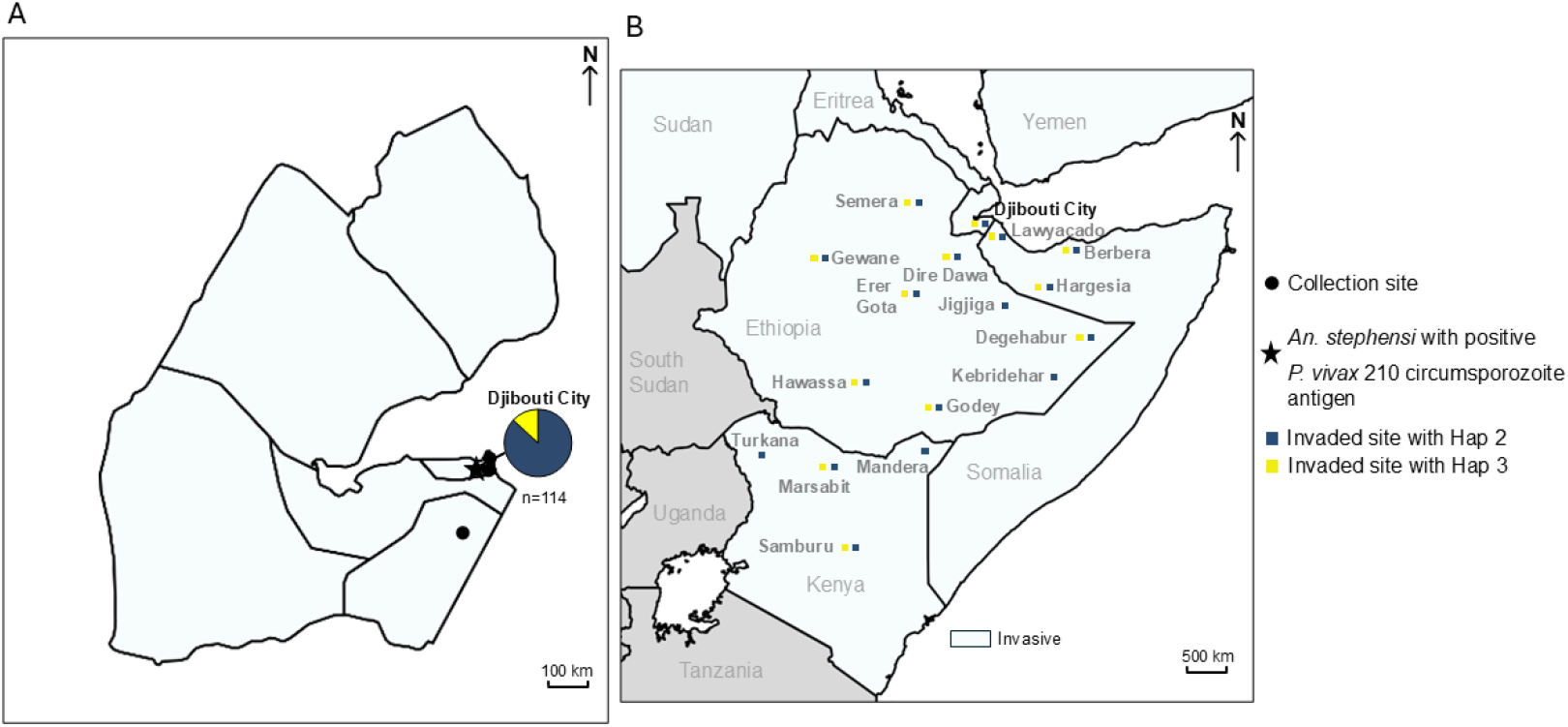
(**A**) Map of collection sites, *Pv*210-positive Djiboutian *An. stephensi* site and COI haplotypes. (**B**) Map of *An. stephensi* COI haplotype 2 and haplotype 3 distribution across the Horn of Africa. Horn of Africa map includes COI data from previously published reports (Appendix). Maps were created by using MapChart (https://www.mapchart.net).

The detection of *Pv*210 circumsporozoite protein using the most advanced csELISA approach in a Djiboutian *An. stephensi* population provides evidence for a direct link between *An. stephensi* and malaria transmission in Djibouti. Thus, our finding supports the previous report of *P. vivax* detection in *An. stephensi* in Djibouti City (9) and suggests that *An. stephensi* may potentially be driving the high incidence rates of *P. vivax* malaria observed in Djibouti in addition to *P. falciparum* malaria (1, 23) (Figure 2). The fact that these malaria-carrying *An. stephensi* were also observed to be closely related to *An. stephensi* populations across the Horn of Africa suggests that the invasive mosquito is potentially contributing to the transmission of malaria in other invaded areas across the region as well (Figure 1).

**Figure 2.**
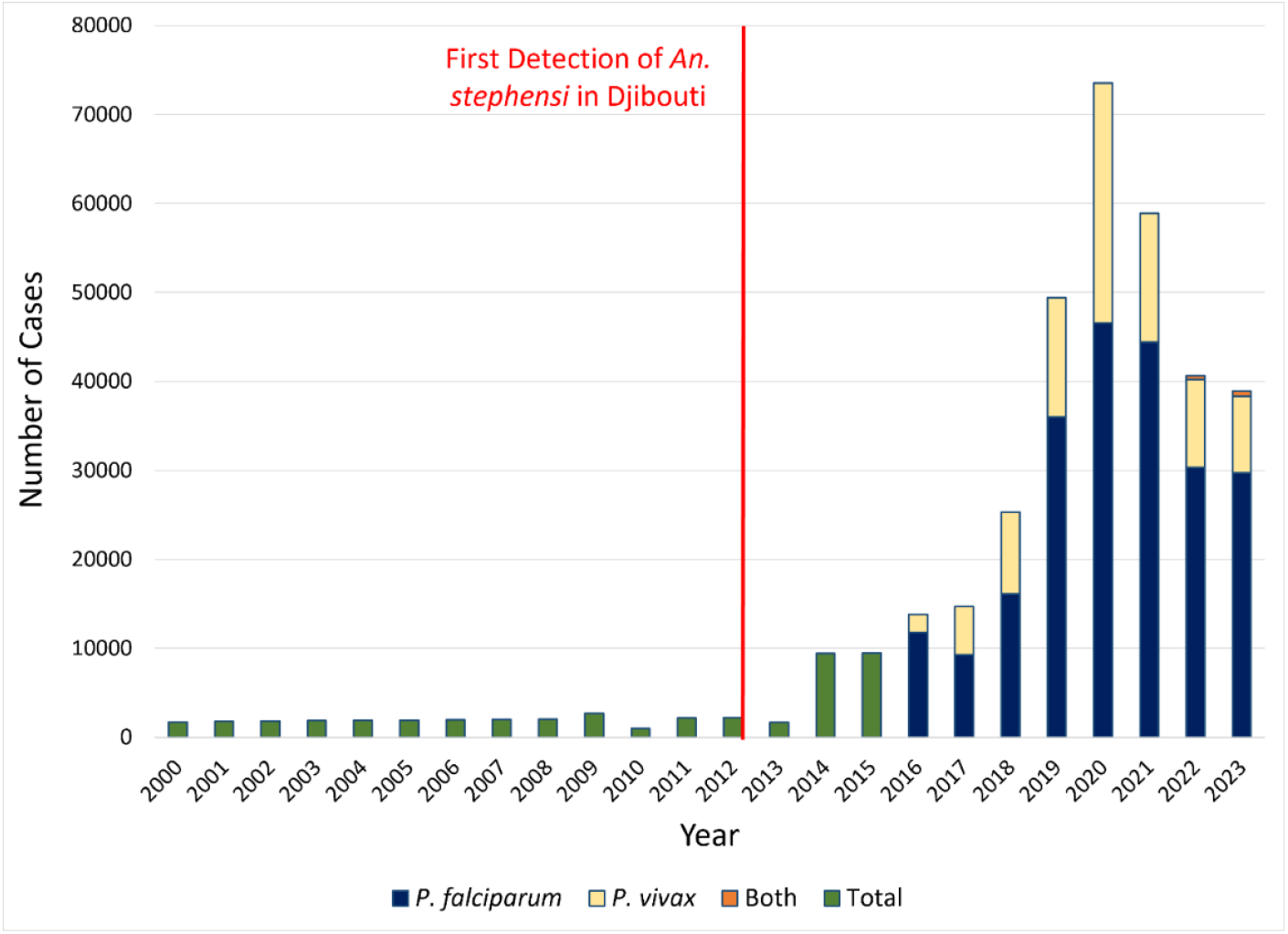
Yearly cases of malaria in Djibouti, 2000-2023. Determination of the *Plasmodium* species and strains began in 2016. Data from World Malaria Reports (2023, 2024) (1, 23).

Taken together, our findings fill critical gaps by (1) providing information on the detection of circumsporozoite-positive *An. stephensi* in Djibouti given that the last detection occurred seven years ago (9), (2) by providing the first molecularly confirmed circumsporozoite-positive *An. stephensi* using the innovative multiplex bead-based csELISA approach (12), and (3) by characterizing the population structure of a recent *An. stephensi* population in Djibouti and its relatedness to other *An. stephensi* populations in the Horn of Africa. This highlights the importance of incorporating both the conventional sporozoite testing using the gold-standard csELISA and population genetics into routine molecular surveillance to better understand the implication of malaria vectors in the disease transmission dynamics, especially in an invasion setting, to inform control strategies.

In Djibouti, the conventional vector control measures will need adjustments to account for *An. stephensi* distinct behaviors as, contrary to the native African malaria vectors, they bite humans both indoors and outdoors and breed in artificial and natural environments (5, 24). Through these behaviors, *An. stephensi* can reach a greater number of human hosts and occupy a greater geographical range than previously prevalent African malaria vectors (24, 25). With *An. stephensi* displaying outdoor biting behaviors, the use of indoor residual spraying (IRS) and insecticide-treated nets (ITNs) alone is insufficient to protect people from bites (26).

Furthermore, *An. stephensi* is displaying resistance to several insecticide classes, including pyrethroids, the insecticide most commonly used for IRS and ITNs (26, 27). This makes control through IRS and ITNs more challenging. Additionally, Djibouti’s coastal location contributes to frequent rainfall and flooding, which increases the number of adult mosquitoes and can increase larval habitats (8, 28). Larval source management (LSM) combats this by either killing larvae or modifying their habitats to contain the spread of populations (29). However, LSM does not provide information on transmission indicators, as the circumsporozoite protein can only be detected in adult mosquitoes. Thus, based on the limitations of IRS, ITNs, and LSM, methods that track entomological indicators for disease transmission, such as increasing adult surveillance, would be beneficial for Djibouti.

## Conclusion

Through this study, we molecularly confirmed for the first time that invasive *An. stephensi* is transmitting malaria parasites (*P. vivax*) in Djibouti. We additionally confirmed through a COI region analysis that *An. stephensi* populations in Djibouti are closely related to the majority of *An. stephensi* populations found in the Horn of Africa. We found that the *An. stephensi* of Djibouti are a likely contributor and possible driver of the transmission of malaria through the Horn of Africa. These findings highlight the threat of *An. stephensi* invasion and support a rapid and comprehensive response to mitigate the harm that *An. stephensi* populations cause, particularly through surveillance and control of adult populations.

## Declarations

### Ethics Approval and Consent to Participate

Not applicable

### Consent for Publication

Not applicable

## Availability of Data and Materials

The sequences generated in this study are available through NCBI Genbank Accession numbers PX682928 – PX683039.

## Funding

Subhika Rao is funded through Public Health Entomology for All (PHEFA) a partnership with the Entomological Society of America (ESA) and Centers for Disease Control and Prevention (CDC).

### Authors’ contributions

SR, JNS, CR, PM, AAA, and SZ contributed to the conception and design of the project. PM, SC, MB, BAK, SKG, MMI, AAA, and SZ organized samples collection. SR, JNS, and CR generated and analyzed the data. SR, JNS, CR, and SZ wrote the first draft of the manuscript. All authors reviewed and approved the final manuscript.

## Acknowledgements

We would like to thank Alice Sutcliffe for discussions on sporozoite detection thresholds using the bead assay method, and the Malaria Research Reference Reagent Resource Center (MR4) and BEI Resources. We would also like to thank the National Malaria Program in Djibouti for coordination of this project.

## Disclaimer

The findings and conclusions expressed herein are those of the author(s) and do not necessarily represent the official position of the US Centers for Disease Control and Prevention.

## Supplemental Information

See attached.

## Reference

1. World Malaria Report [Internet]. 2024. Available from: https://www.who.int/teams/global-malaria-programme/reports/world-malaria-report-2024.

2. Faulde MK, Rueda LM, Khaireh BA. First record of the Asian malaria vector Anopheles stephensi and its possible role in the resurgence of malaria in Djibouti, Horn of Africa. Acta Trop. 2014;139:39–43.

3. Malaria Threats Map [Internet]. 2025. Available from: https://apps.who.int/malaria/maps/threats/.

4. Ollivier L, Nevin RL, Darar HY, Bougère J, Saleh M, Gidenne S, et al. Malaria in the Republic of Djibouti, 1998-2009. Am J Trop Med Hyg. 2011;85(3):554–9.

5. Samake JN, Yared S, Hassen MA, Zohdy S, Carter TE. Insecticide resistance and population structure of the invasive malaria vector, Anopheles stephensi, from Fiq, Ethiopia. Sci Rep. 2024;14(1):27516.

6. Yared S, Gebresilassie A, Aklilu E, Abdulahi E, Kirstein OD, Gonzalez-Olvera G, et al. Building the vector in: construction practices and the invasion and persistence of Anopheles stephensi in Jigjiga, Ethiopia. Lancet Planet Health. 2023;7(12):e999–e1005.

7. de Santi VP, Khaireh BA, Chiniard T, Pradines B, Taudon N, Larréché S, et al. Role of Anopheles stephensi Mosquitoes in Malaria Outbreak, Djibouti, 2019. Emerg Infect Dis. 2021;27(6):1697–700.

8. Zayed A, Moustafa M, Tageldin R, Harwood JF. Effects of Seasonal Conditions on Abundance of Malaria Vector Anopheles stephensi Mosquitoes, Djibouti, 2018-2021. Emerg Infect Dis. 2023;29(4):801–5.

9. Seyfarth M, Khaireh BA, Abdi AA, Bouh SM, Faulde MK. Five years following first detection of Anopheles stephensi (Diptera: Culicidae) in Djibouti, Horn of Africa: populations established-malaria emerging. Parasitol Res. 2019;118(3):725–32.

10. Bangs MJ, Rusmiarto S, Gionar YR, Chan AS, Dave K, Ryan JR. Evaluation of a dipstick malaria sporozoite panel assay for detection of naturally infected mosquitoes. J Med Entomol. 2002;39(2):324–30.

11. Emiru T, Getachew D, Murphy M, Sedda L, Ejigu LA, Bulto MG, et al. Evidence for a role of Anopheles stephensi in the spread of drug- and diagnosis-resistant malaria in Africa. Nat Med. 2023;29(12):3203–11.

12. Sutcliffe AC, Irish SR, Rogier E, Finney M, Zohdy S, Dotson EM. Adaptation of ELISA detection of Plasmodium falciparum and Plasmodium vivax circumsporozoite proteins in mosquitoes to a multiplex bead-based immunoassay. Malar J. 2021;20(1):377.

13. Ismael Dini M, Abdoul-latif F, Ismael Y, Elmi Fourreh A, Ali A, Waiss I, et al. Malaria epidemiology in Djibouti (2019-2024) and predictive modeling for 2025 using computational approaches. 2024.

14. Waymire E, Samake JN, Gunarathna I, Carter TE. A decade of invasive Anopheles stephensi sequence-based identification: toward a global standard. Trends Parasitol. 2024;40(6):477–86.

15. Rozas J, Ferrer-Mata A, Sánchez-DelBarrio JC, Guirao-Rico S, Librado P, Ramos-Onsins SE, et al. DnaSP 6: DNA Sequence Polymorphism Analysis of Large Data Sets. Mol Biol Evol. 2017;34(12):3299–302.

16. Stamatakis A. RAxML version 8: a tool for phylogenetic analysis and post-analysis of large phylogenies. Bioinformatics. 2014;30(9):1312–3.

17. Rambaut A. Figtree ver 1.4.4. Institute of Evolutionary Biology, University of Edinburgh, Edinburgh. 2018 [Available from: https://tree.bio.ed.ac.uk/software/figtree/.

18. Carter TE, Yared S, Getachew D, Spear J, Choi SH, Samake JN, et al. Genetic diversity of Anopheles stephensi in Ethiopia provides insight into patterns of spread. Parasit Vectors. 2021;14(1):602.

19. Hawaria D, Kibret S, Zhong D, Lee M-C, Lelisa K, Bekele B, et al. First report of Anopheles stephensi from southern Ethiopia. Malaria Journal. 2023;22(1):373.

20. Ali S, Samake JN, Spear J, Carter TE. Morphological identification and genetic characterization of Anopheles stephensi in Somaliland. Parasit Vectors. 2022;15(1):247.

21. Samake JN, Athinya DK, Milanoi S, Ramaita E, Muchoki M, Omondi S, et al. Spatial distribution and population structure of the invasive Anopheles stephensi in Kenya from 2022 to 2024. Sci Rep. 2025;15(1):19878.

22. Samake JN, Lavretsky P, Gunarathna I, Follis M, Brown JI, Ali S, et al. Population genomic analyses reveal population structure and major hubs of invasive Anopheles stephensi in the Horn of Africa. Mol Ecol. 2023;32(21):5695–708.

23. World Malaria Report [Internet]. 2023. Available from: https://www.who.int/teams/global-malaria-programme/reports/world-malaria-report-2023.

24. Mwema T, Zohdy S, Sundaram M, Lepczyk CA, Narine L, Willoughby JR. A quantitative and systematic analysis of Anopheles stephensi bionomics and control approaches. Acta Trop. 2024;260:107431.

25. Sinka ME, Pironon S, Massey NC, Longbottom J, Hemingway J, Moyes CL, et al. A new malaria vector in Africa: Predicting the expansion range of Anopheles stephensi and identifying the urban populations at risk. Proc Natl Acad Sci U S A. 2020;117(40):24900–8.

26. Mnzava A, Monroe AC, Okumu F. Anopheles stephensi in Africa requires a more integrated response. Malar J. 2022;21(1):156.

27. Yared S, Gebressielasie A, Damodaran L, Bonnell V, Lopez K, Janies D, et al. Insecticide resistance in Anopheles stephensi in Somali Region, eastern Ethiopia. Malar J. 2020;19(1):180.

28. Imbahale SS, Paaijmans KP, Mukabana WR, van Lammeren R, Githeko AK, Takken W. A longitudinal study on Anopheles mosquito larval abundance in distinct geographical and environmental settings in western Kenya. Malar J. 2011;10:81.

29. Okumu F, Moore SJ, Selvaraj P, Yafin AH, Juma EO, Shirima GG, et al. Elevating larval source management as a key strategy for controlling malaria and other vector-borne diseases in Africa. Parasit Vectors. 2025;18(1):45.

