## Supplemental Figure S1 for "Evidence of *Anopheles stephensi* involvement in the transmission of *Plasmodium vivax* in Djibouti, 2024"

### Supplementary Materials

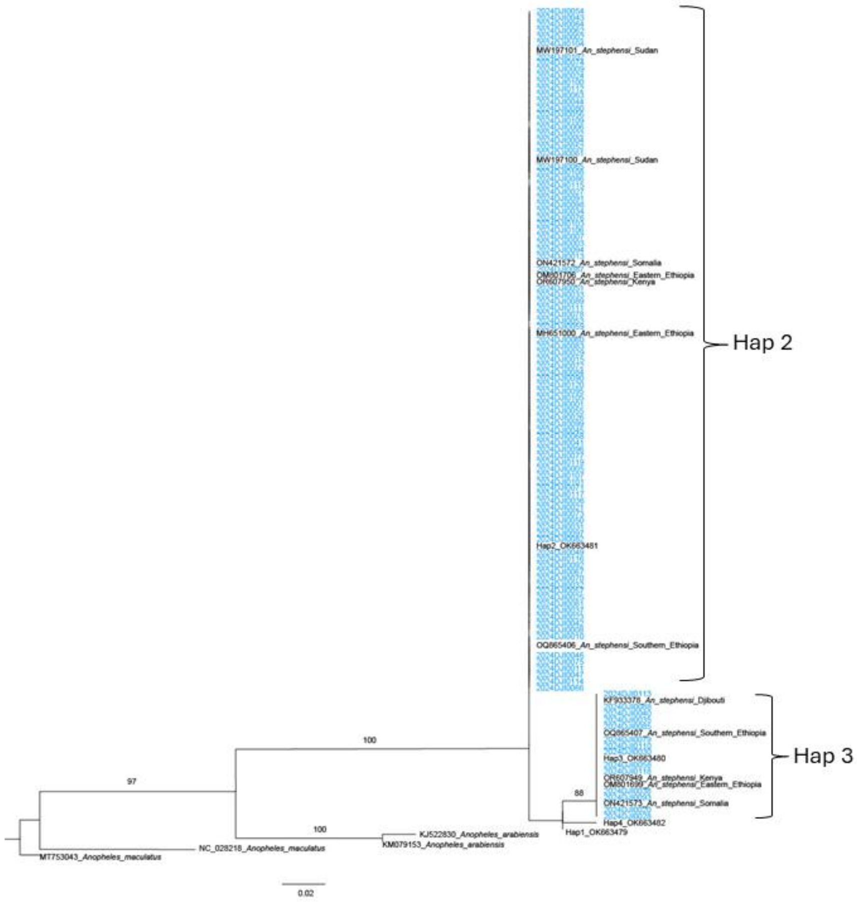


**Figure S1.** Phylogenetic tree of Horn of Africa *An. stephensi* cytochrome oxidase subunit I (COI). The evolutionary history was inferred by using the Maximum Likelihood method based on the General Time Reversible model. Blue = Djibouti COI data generated in this study. Numbers on the branches indicate bootstrap values (only values > 70 are shown). Scale bar indicates the number of nucleotide substitutions per site. Haplotype numbering was based on haplotypes list from Carter et al. 2021(1).

**Reference**

1. Carter TE, Yared S, Getachew D, Spear J, Choi SH, Samake JN, et al. Genetic diversity of *Anopheles stephensi* in Ethiopia provides insight into patterns of spread. Parasit Vectors. 2021;14(1):602.
